# Self-Assembled siRNA-Gold Supraclusters Detected at the Single-Molecule Level in the NIR-II Window

**DOI:** 10.1101/2025.06.01.657319

**Authors:** Zeineb Ayed, Blanca Martín Muñoz, Catharina I. Vandekerckhove, Virginie Faure, Syed Nafis Shadman Ali, Jean-luc Coll, Laurent Cognet, Xavier Le Guével

**Affiliations:** University Grenoble Alpes, INSERM U1209, CNRS UMR5309, Institute for Advanced Biosciences F-38000 Grenoble, France; Univ. Bordeaux, Laboratoire Photonique Numérique et Nanosciences (LP2N), UMR 5298, F-33400 Talence, France; Institut d’Optique & CNRS, LP2N UMR 5298, F-33400 Talence, France

**Author notes:** **Corresponding Author** Xavier Le Guével;, Laurent Cognet.

**Keywords:** gold nanoclusters, fluorescence, assembly, single-particle tracking

## Abstract

Gold nanoclusters (AuNCs) possess unique photophysical properties that make them excellent candidates for advanced bioimaging and single-particle detection. In this work, we report the self-assembly of highly emissive, positively charged NIR-II AuNCs stabilized by cysteamine, directed by small interfering RNA (siRNA), which serves as both a structural and electrostatic modulator. The resulting supramolecular assemblies exhibit quasi-spherical morphologies around 100 nm in diameter, with outstanding colloidal stability, photostability, and enzymatic resistance. Their strong photoluminescence, extending up to 1400 nm, enables robust single-particle detection in solution. Spectroscopic and structural analyses—including fluorescence spectroscopy, small-angle X-ray scattering (SAXS), and single-particle tracking—highlight the pivotal role of siRNA in tuning the assembly process via charge balance and concentration- dependent interactions. Beyond providing insights into the structural and photophysical behavior of nucleic acid–guided AuNC assemblies, these results underscore their promise as multifunctional nanoplatforms for integrated imaging and gene-silencing therapies in biophotonic and theranostic applications.

---

Gold nanoclusters (AuNCs) represent an emerging class of luminescent nanomolecular species with unique molecular-like-properties that has been exploited for several applications in optoelectronics^1^, photocatalysis^2^, bioimaging^3, 4^, biosensing^4, 5^, and therapy^6^. Recently, several groups have introduced approaches for producing assembled nanostructures with AuNCs by physical interaction^7^ or with the use of different matrices (small organic molecules^8^, polymers^9^, MOF^10^, nucleic acids^11-13^) which outperform individual AuNC photonic properties such as improved brightness^14^, prolonged fluorescence lifetime^15^, or enhanced two-photon absorption enhancement^16^. Excellent reviews reported the advances in this field^17^. In addition, such self-assembled nanostructures were proposed as theranostic agents and offer the possibility to be activated by X-ray or near infrared light and to deliver therapeutic agents or biological targets. For instance, us and others have demonstrated how self-assembled AuNCs–siRNA can target telomeric response for radiotherapy enhancement^11^ or to enhance the anti-tumor immune response of stereotactic ablative radiotherapy at primary and metastatic tumors^12^.

In parallel, there are lot of efforts to develop nanoparticles or nanostructures that can be detected and tracked at the single particle level to probe and decipher biological structures at the cellular level^18^. While several candidates have shown outstanding spatial and temporal resolution in the visible (400-700 nm) and in the first infrared window (700-900 nm), only few biocompatible candidates such as carbon nanotubes^19^ and highly loaded polymeric nanoparticles^20^ have been detected at the single particle level in the second biological window, called NIR-II or SWIR (shortwave infrared) between 900-1700 nm,. NIR-II region is particularly appealing for biological and biomedical studies due to the reduced autofluorescence and scattering which should improve the resolution in depth and find some interest in complex cellular environments (organoids, organ-on-chip) and in living tissues^21^. Most existing NIR II probes possess imaging capabilities but lack therapeutic functionalities. Therefore, new probes combining both NIR-II detection and therapeutic modalities would be of high interest for bio(medical)applications.

In this work, we demonstrate that ligand engineering can enhance the quantum yield of positively charged NIR-II-emitting AuNCs. Furthermore, we show that, in the presence of siRNA, these AuNCs can self-assemble into quasi-spherical structures around 100 nm in size via electrostatic interactions. For this study, we used siHSF1 (silencing heat shock transcription factor 1)^22^ as a model siRNA. These AuNCs–siRNA supraclusters demonstrate an excellent photo and enzymatic stabilities, an enhanced photoluminescence (PL) shifted to longer wavelengths compared to single AuNCs, and are detectable at the single particle level above 1000 nm wavelengths.

**Scheme 1.**
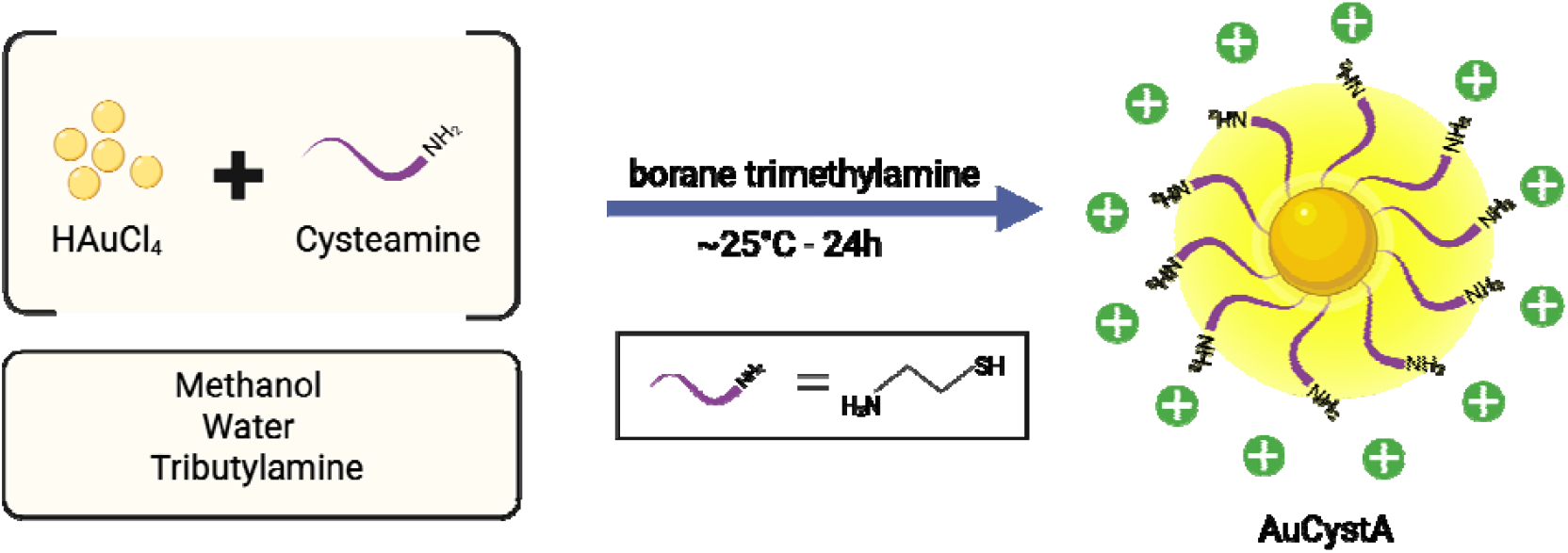
Schematic illustration of the synthesis of cysteamine stabilized gold nanoclusters denoted AuCystA.

Positively charged gold nanoclusters (AuCystA) were synthesized using cysteamine as a capping ligand and Me□NBH□as a mild reducing agent (**Scheme 1, see experimental section in SI)**. Transmission Electron Microscopy (TEM) images reveal an average core for diameter of 2–4 nm with lattice fringes spaced at approximately 0.23 nm, characteristic of the Au(111) plane (**Figure 1a**). Mass spectrometry (MALDI-ToF) further identify two predominant species (**Figure 1b**) with two main broad peaks (m/z = 5700 and m/z = 8600), most likely corresponding to Au□ □CystA□ □and Au□ □CystA□ □ respectively. Zeta potential measurements confirm the positive charge of AuCystA in a wide pH range (4-9) over at least a week due to the primary amine of cysteamine (**Figure 1c, Figure S1**).

**Figure 1.**
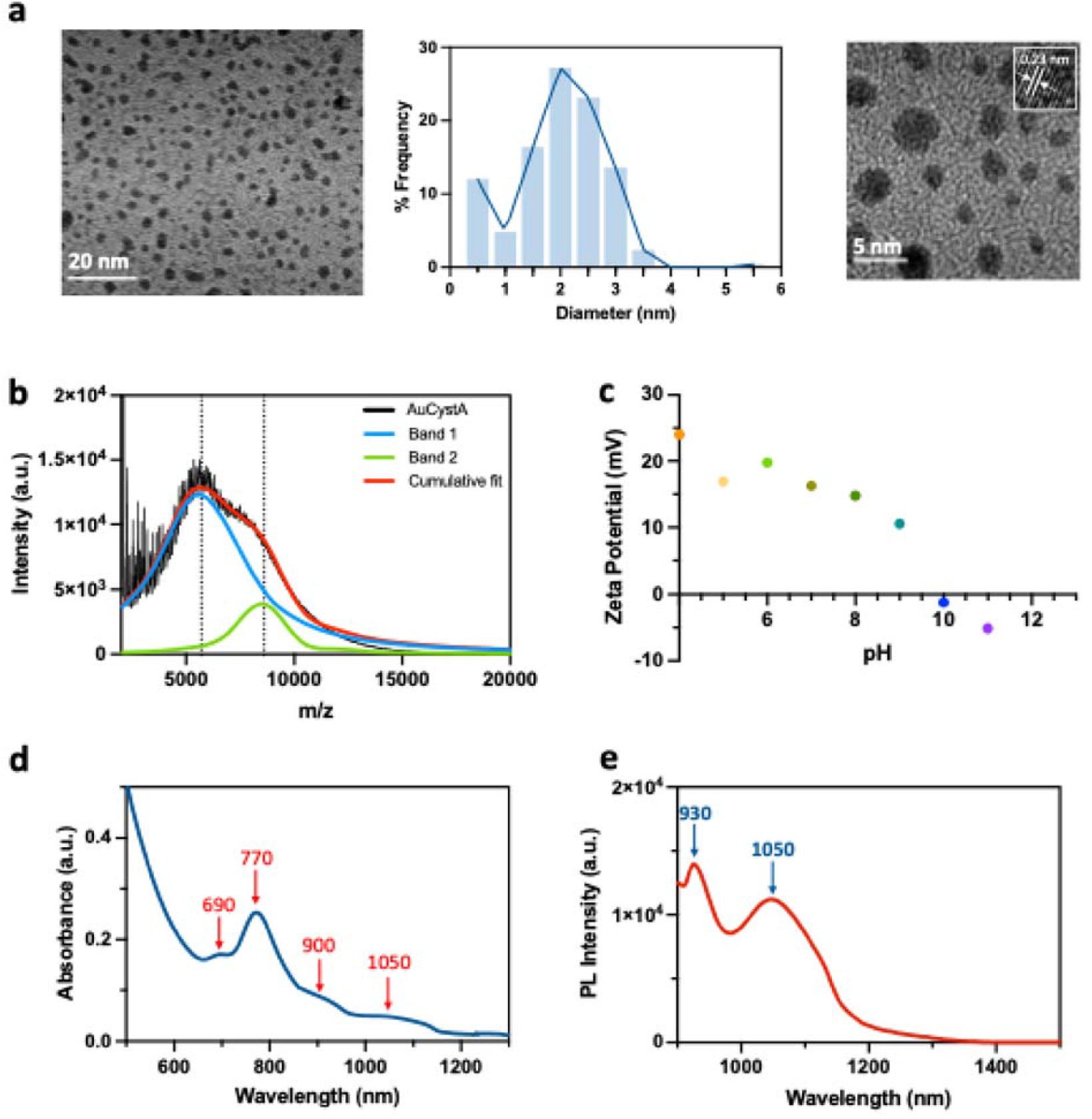
**(a)** HRTEM (High-Resolution Transmission Electron Microscopy) images of AuCystA and histogram of the average size determined from HRTEM images using 250 particles. Zoomed TEM image with inset showing the distance between two lattice fringes (∼___0.23 nm: (Au 111)). **(b)** MALDI-Tof spectrometry of AuCystA nanoclusters and deconvolution of the bands 1 (m/z = 5700, Au_25_CystA_18_) and 2 (m/z = 8600, Au_36_CystA_24_). **(c)** Zeta potential of AuCystA in PBS at different pH. **(d)** Absorbance spectrum of AuCystA in water. **(e)** PL spectrum of AuCystA in water. A_exc._ 808 nm.

Absorbance spectrum of AuCystA reveal distinct absorption bands at 690, 770, 900, and 1050 nm (**Figure 1d**). The absorption band at 690 nm is commonly associated with the Au_25_ structure^23^. Interestingly, the near-infrared bands observed in AuCystA resemble those found in AuNCs stabilized by co-ligands, where anisotropic surface charges induce structural distortions at varying distances^24, 25^. However, in the present case, cysteamine alone stabilizes these nanoclusters. One possible explanation for the presence of the near infrared bands could be related to the spatial arrangement of the cysteamine on AuNCs surface^25^. Alternatively, the bands may result from excited-state exciton localization, induced by structural distortions within the gold core of the second species, AuLLCystALL. Indeed, this is consistent with previous reports on rod-like AuNCs with comparable atomic numbers, such as [AuLL(PPhL)LL(SCLHLPh)LLXL]L (X = Cl/Br)^26^ and Au_38_SG_24_^27^ which exhibit similar absorption bands at 800 and 1000 nm.

Upon excitation at 808 nm, AuCystA exhibits strong photoluminescence (PL) in the NIR-II region, characterized by a prominent emission band at 930 nm and a second PL emission band centred at 1050 nm (**Figure 1e**). This dual-emission feature suggests the presence of two distinct emissive states or pathways within the nanocluster^28^. Additional PL measurements at different excitation wavelengths (400 nm, 690 nm, and 780 nm) yield consistent emission spectra, (**Figure 1e, Figure S2a**) and confirm the presence of the excitation/emission coupling at 770 nm/1050 nm (**Figure S2b)**. We estimated the photoluminescence quantum yield (PLQY) of AuCystA is at 7.1% using indocyanine green (ICG, PLQY 0.23%) as a reference which is more than 14 times higher than the well-described water soluble Au_25_SG_18_ (SG : glutathione)^29^. Furthermore, time-resolved PL analysis of the 1050 nm emission band, fitted using a biexponential decay model, reveal two distinct emissive processes: a fast decay component with a lifetime of ∼48 ns and a slower decay component of ∼1.2 μs (**Figure S3**). The shorter lifetime (τ_1_ = 48 ns) likely corresponds to rapid radiative transitions associated with the gold core’s intrinsic electronic states, whereas the longer lifetime (τ_2_ = 1.2 μs) suggests additional relaxation pathways, potentially involving surface states modulated by cysteamine ligands.

While PL of AuCystA is sensitive to both pH and solvent polarity (**Figures S4a-b**), at pH > 9 the PL intensity decreases, likely due to the deprotonation of the cysteamine ligand. Conversely, at mildly acidic pH (4–5), the PL intensity increases, suggesting that ligand protonation stabilizes the emissive states—a trend previously reported for analogous gold nanocluster systems^30^. In polar solvents, such as dimethyl sulfoxide (DMSO) and ethanol, AuCystA maintains strong PL, whereas in acetonitrile, PL is quenched despite an unchanged absorbance spectrum (**Figures S4c-d**). This suggests that the solvent interactions or polarity effects disrupt excited-state processes, further confirming the important role of ligand protonation state in the PL properties of AuCystA.

Additional studies assessed the stability of AuCystA’s PL in various biological buffers confirming robust fluorescence in BSA, HBSS at both pH 7 and pH 5, and 0.9% NaCl solution (**Figure S5**). This stability highlights the potential of AuCystA nanoclusters for *in vitro* and *in vivo* applications where bright PL in complex biological media would be essential for monitoring^31^. Given the strong influence of pH and solvent polarity on AuCystA’s PL, we further explored their optical properties in water (HLO) and deuterium oxide (DLO). The D_2_O experiment shows an enhancement of the band at 1050 nm when exciting at 780 nm suggesting energy transfer from AuCystA surface. Excitations at 450 nm and 690 nm show an overall increase of the full PL spectra indicating more complex energy transfer taking place in the core and on the surface (**Figure S6**)^32^.

The NIR-II detection sensitivity of AuCystA was next evaluated. First, we assessed its limit of detection in a capillary tube (1Lmg/mL) immersed in 1% Intralipid^®^, where the signal remained detectable up to a depth of 1.4Lcm (**Figure S7**). Second, AuCystA was detectable in both water and plasma across various sub-NIR-II windows at concentrations as low as 0.1Lμg/mL (∼12LnM), highlighting their exceptional sensitivity and robust detection performance in complex biological media (**Figure S8**).

In the following, the positively charged AuCystA nanoclusters were mixed with the negatively charged siRNA (**see SI)**, leading to spontaneous self-assembly of AuCystA–siRNA structures driven by electrostatic interactions (**Figure 2a**). To assess the impact of different ratios on complex formation, an agarose gel shift assay was performed with various AuCystA:siRNA weight ratios (1:1, 2:1, 5:1, and 9:1; **Figure 2b**). As expected, free AuCystA nanoclusters accumulate at the top of the gel near the negative electrode, while free siRNA migrates toward the positive electrode. At a 1:1 and 2:1 ratio, a significant portion of siRNA remains unbound, as evidenced by its distinct migration pattern with a rather weak PL signal of these assemblies. Increasing the weight ratio to 5:1 or 9:1 leads to complete binding of siRNA to AuCystA. The zeta potential of the assembled AuCystA–siRNA at the weight ratio 5:1 is slightly negative (−7±1 mV) compare to the free siRNA (−30mV<), confirming the full complexation of the siRNA.

**Figure 2.**
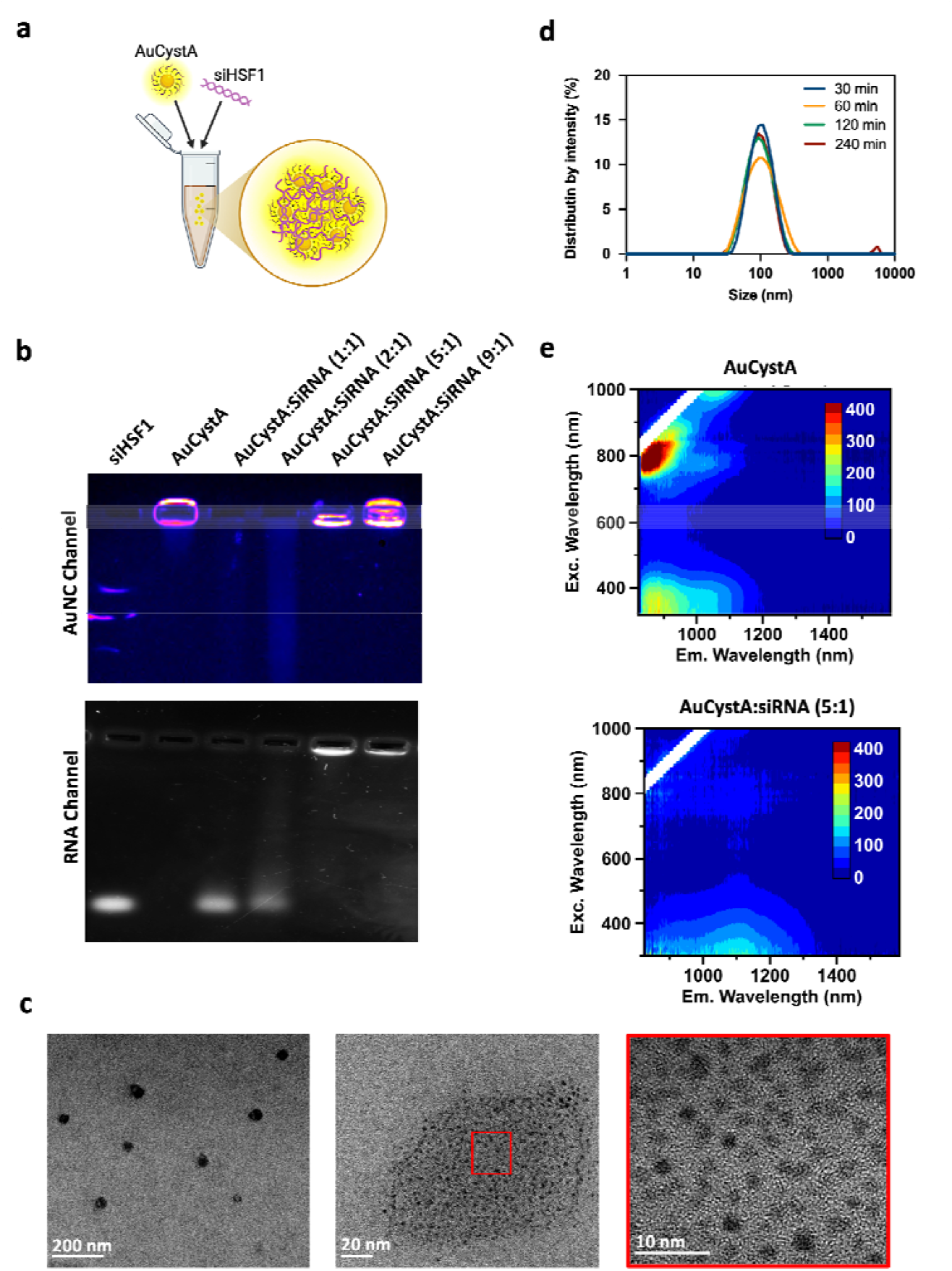
**(a)** Schematic representation of self-assembled AuNCs. **(b)** Migration of AuCystA-siRNA by electrophoresis in 1% agarose gel using the fluorescent channel of AuNCs (up) and after siRNA staining with GoldSyber (down) at different AuCystA:siRNA weight ratio. **(c)** TEM images of the self-assembled AuCystA:siRNA (weight ratio 5:1) at different magnifications. **(d)** Hydrodynamic diameters of the self-assembled AuCystA:siRNA (weigth ratio 5:1) overtime. **(e)** Excitation-photoluminescent maps of the AuCystA (up) and the self-assembled AuC-siRNA (down).

For the remainder of the study, we used a 5:1 ratio as the optimal condition. When other ratios were used, they were explicitly indicated in the complex name. TEM measurements show predominantly spherical assemblies ranging from 30 to 100 nm in diameter with individual AuCystA nanoclusters visible evenly distributed dark spots less than 3 nm within the assemblies for the 5:1 ratio (**Figure 2c, S9)**. The presence of gold in the supraclusters is also confirmed by EDS (**Figure S10**). In contrast, only dispersed AuCystA are observed at the ratio 1: 1 (**Figure S9**).

The resulting complex AuCystA–siRNA was characterized using dynamic light scattering (DLS) indicating an average hydrodynamic diameter of approximately 100 nm (**Figure 2d**), which remains stable over a period of at least 240 minutes. Complementary SAXS measurements yielded an average radius of 65 nm and a slight shift in the size curve over time suggesting that the assemblies might become progressively more compact (**Figure S11**).

The protective efficacy of AuCystA complexation against RNase-mediated degradation of siRNA was then evaluated. Since RNases are abundant in biological environments and rapidly degrade siRNA, we first complexed siRNA with AuCystA at an optimized 1:5 weight ratio. The resulting complexes were then exposed to increasing concentrations of RNase A (1–10 ng/µL), exceeding typical levels found in the bloodstream^6^. Gel electrophoresis reveals that free siRNA is completely degraded at an RNase A concentration of 1 ng/µL. In stark contrast, siRNA complexed with AuCystA remains intact across all tested RNase concentrations (**Figure S12**).

The optical properties of the AuCystA nanocluster and its siHSF1 complex were first characterized in solution using absorbance and PL spectroscopy (**Figure S13**). The absorbance spectra reveals that free siRNA exhibits the typical nucleic acid absorption peak at 260 nm. Upon complexation, the resulting spectrum is essentially an additive overlay of the individual spectra of AuCystA and siRNA, with no significant alteration in shape. Characteristic AuCystA bands at 690 and 770 nm are not observed, likely due to the detection limits of the instrumentation used. The PL spectra of free AuCystA exhibit a broad emission profile with a prominent peak at 1050 nm. Upon introduction of siRNA, a marked decrease in PL intensity is observed, acco by a significant redshift in the emission maximum to approximately 1400 nm. These findings are in agreement with previous results correlating the increase of rigidity in the local environment of AuNCs to a bathochromic of his PL emission spectra^3^. An excitation-photoluminescence map of AuCystA versus AuCystA- siRNA confirms also the shift and decrease intensity observed in the PL spectra of the components when assemblies are formed (**Figure 2e**).

The photostability of free AuCystA nanoclusters and their corresponding AuCystA– siRNA assemblies were also compared to the organic dye ICG (**Figure S14**). Under continuous laser excitation for 20 minutes (120 mW/cm^2^), ICG exhibits a significant loss in intensity—more than 70%—indicating rapid photobleaching. In contrast, free AuCystA nanoclusters demonstrates about a 40% reduction in intensity and less than a 10% drop for AuCystA– siRNA complex. This substantial improvement in photostability upon complexation with siHSF1 suggests that siRNA plays a critical role in protecting AuCystA emission, probably shielding the nanocluster from photodegradation.

To gain a deeper understanding of the formation and properties of the supraclusters, both AuCystA and their corresponding siRNA self-assemblies were studied at the single-particle level. For this, we used a widefield epifluorescence single molecule microscope (**Figure 3a**), equipped with 488 nm laser (1.4 kW/cm^2^) and an InGaAs camera, which has excellent sensitivity in the NIR-II window. **Figure 3b** shows representative PL images of solutions containing AuCystA– siRNA complexes when different weight ratios of AuCystA: siRNA were used. As a control, we first imaged free AuCystA in the absence of siRNA, observing the formation of a homogeneous luminescent layer on the glass slide. In contrast, when siRNA was present, increasing the concentration of AuCystA led to the formation of self-assembled particles composed of AuCystA clusters bound to siRNA, which appeared as bright particles in suspension. The optimal ratio for stable self-assembly of the AuCystA– siRNA complex was determined to be 5:1, as evidenced by the presence of uniform, spherical, and diffusive structures corresponding to complete siRNA binding (**Figure 2b**). Zeta potential of AuCystA– siRNA 5:1 is almost neutral (−7±5 mV due to the full complexation of siRNA and the presence of AuCystA. At sub-optimal weight ratios such as 2:1, only dispersed AuCystA was observed, whereas exceeding the optimal range (e.g., at a 9:1 weight ratio) resulted in heterogeneous, large aggregates lacking structural control (**Figure S15**).

**Figure 3.**
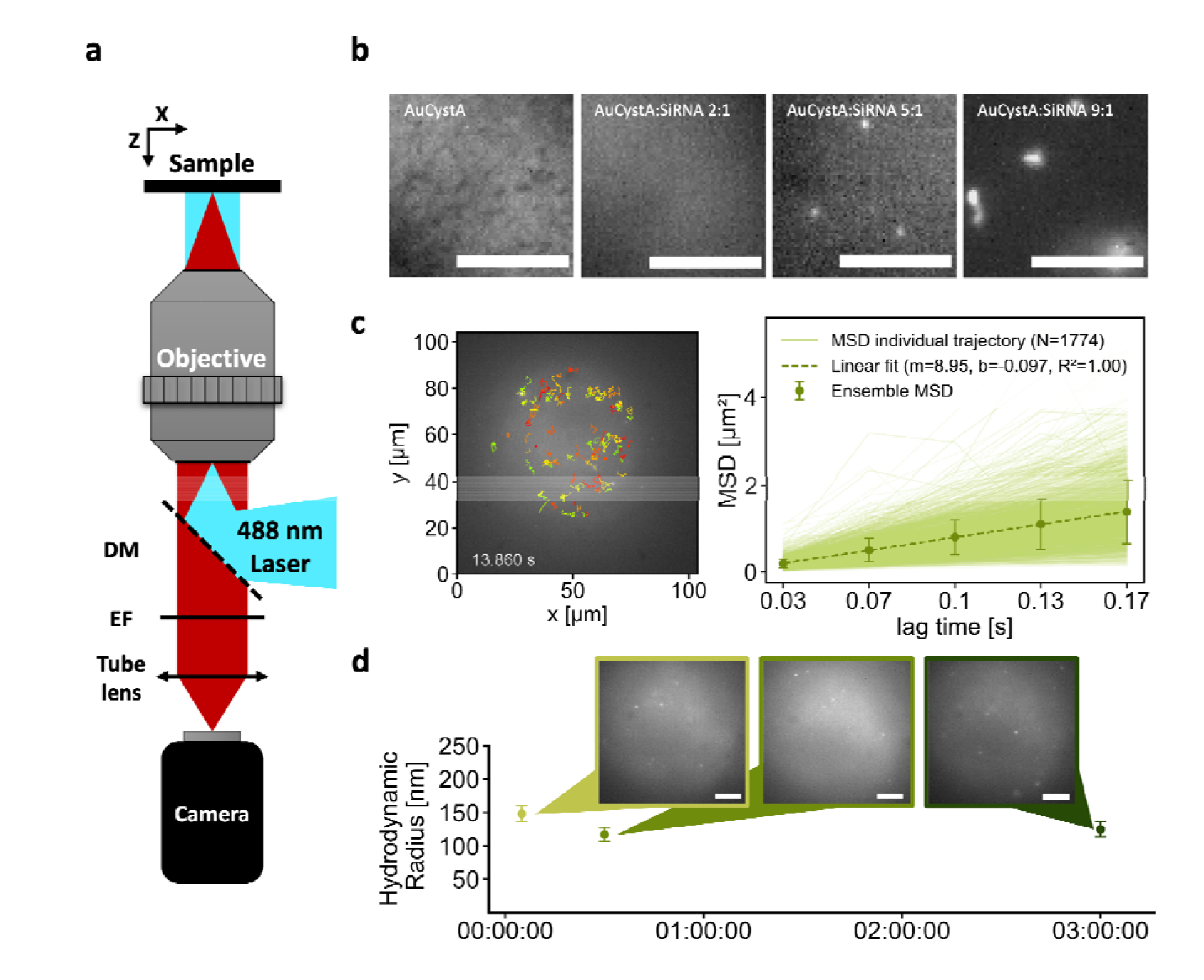
NIR II microscopy detection as a function of complexation. **a**. Schematic of the experimental setup: DM-dichroic mirror, EF-emission filter. **b**. Complexation at different concentrations. Sample frames are shown for each case. From left to right: control (AuCystA without siRNA), AuCystA:siRNA at 2:1 weight ratio (stabilizatio phase), AuCystA:siRNA 5:1 (assembly formation), AuCystA:siRNA 9:1 (formation of heterogeneous structures). Scale bar = 10 µm. **c**. MSD analysis of AuCystA-siRNA assemblies (5:1). Left: Example frame showing detected particles trajectories. Right: Individual MSDs (light green) from 1774 trajectories across five fields of view in the same sample; ensemble MSD calculated from individual ones (dark green); linear fit over ensemble MSD (dar green dashed line) used to obtain the typical hydrodynamic radius of the tracked objects. **d**. Stability kinetics of AuCystA-siRNA assemblies (5:1). Hydrodynamic radius remains consistent over time (5 minutes, 30 minutes, and 3 hours post-mixing). Sample frames at each time point are shown together with the calculated radius obtained after trajectories analysis. Scale bar = 10 µm.

Next, we analysed the single-particle diffusion properties of these supraclusters to calculate their hydrodynamic radius, in order to provide a quantitative evaluation of their structural characteristics that complement the qualitative insights obtained from microscopy. **Figure 3c** presents an example of such analysis in the specific case of a AuCystA- siRNA at a 5:1 weight ratio 30 minutes post-mixing. On the left panel (**Figure 3c**) a representative image is shown where the reconstructed 2D trajectories of the particles are superimposed, after 13 seconds of recording. Owing to the excellent photostability of the assemblies, trajectories lengths are not limited by photobleaching, but only by the depth-of-focus of the microscope as particles diffuse in 3D. Furthermore, in the case of Brownian motion, each dimension can be analysed independently, allowing us to calculate and analyse the 2D mean square displacement (MSD) for every individual trajectory. The graph presented on the right panel of **Figure 3c** compiles results from N = 1774 trajectories along with the linear fit -as expected for Brownian motion- of their average MSD (**video 1**). From the slope of the fit, we could determine the average particle hydrodynamic diameter via the Stokes–Einstein equation (**see Materials & Methods**). The resulting value,101±9 nm, is consistent with the earlier structural analysis.

Finally, we evaluated the stability of the supraclusters, by monitoring the evolution of the particle hydrodynamic radius over time. The PL images (AuCystA: siRNA at 5:1 weight ratio) reveal consistent spherical assembly structures after 5 minutes, 30 minutes and 3 hours post-mixing (**Figure 3d**) confirmed by the average hydrodynamic radius of 148±12 nm, 117±10 nm, and 125±11 nm, respectively. These results are consistent with previous SAXS measurements (**Figure S11**), which show a similar trend, although indicating a progressively more compact size over time.

To investigate conditions beyond the optimal ratio, we performed the same analysis for the 9:1 AuCystA– siRNA system. Under these conditions, an excess of AuCystA led to the visible formation of large AuCystA– siRNA aggregates (**Figure S15a**). Additionally, a slight redshift was observed in the 2D spectral maps (**Figure S15b**), which was notably smaller than that observed at the 5:1 weight ratio (**Figure 2e**), likely due to the predominance of unbound AuCystA in the solution. Finally, diffusion analysis revealed the presence of large aggregates with a hydrodynamic radius of approximately 1.5 ± 0.15 µm (**Figure S15c, video 2**). Thes findings further support the conclusion that an excessive concentration of AuCystA promotes the formation of uncontrolled, heterogeneous structures.

## CONCLUSION

In this work, we have engineered a positively charged gold nanocluster (AuCystA) that emits brightly in the NIR-II window with a high quantum yield (7.1%) and demonstrate strong stability in biological media. Using the AuCystA scaffold, we formed ∼100 nm supraclusters assemblies with siRNA driven by electrostatic interactions. The assemblies revealed efficient protection of siRNA against RNase-mediated degradation and a strong photostability. Notably, the resulting assemblies exhibit broad, red-shifted NIR-II emission extending up to 1400 nm, with sufficient brightness to enable reliable detection and sizing at the single-particle level. While further work is needed to explore these self-assembled siRNA-Gold supraclusters behaviour in complex biological environments, these results provide a clear proof of concept that such assemblies are both structurally and optically suited to serve as theranostic agents that can be monitored at the single-particle level in the biological tissue transparency window. This study marks a foundational step toward the development of modular, NIR-II active nanoplatforms for imaging and possible targeted delivery in biomedical contexts.

## ASSOCIATED CONTENT

### Supporting Information

Materials & Methods; physico-chemical, optical and morphological characterizations of single and assembled gold nanoclusters are reported in the supporting documents available online.

## AUTHOR INFORMATION

### Corresponding Author

*Xavier Le Guével;

*Laurent Cognet;

### Author Contributions

The manuscript was written through contributions of all authors. All authors have given approval to the final version of the manuscript.

### Funding Sources

X.LG: SIREN “ANR-20-CE92-0039-01”, NAnoGOLD “ANR-22-CE29-0022”) X.LG, JL.C: ANR SEQUOIA “ANR-22-CE18-0006”.

L.C. acknowledges support from CNRS, the University of Bordeaux, the France-BioImaging National Infrastructure (ANR-10-INBS-04-01), the Idex Bordeaux (Grand Research Program GPR LIGHT) and the European Research Council Synergy grant (951294).

L.C. and X.LG acknowledge support from Agence Nationale de la Recherche (ANR-24-CE09- 2351-01).

## ACKNOWLEDGMENT

XLG would like to thank Laetitia Rapenne for the HRTEM measurements, Aurélien Thureau for the SAXS measurements, Thibault Gallavardin and Iyanu Diriwari for the fluorescence lifetime measurements. LC and BMM would like to thank M. Tondusson, Q. Gresil and R. Mangalwedhekar for providing experimental help and support.

## ABBREVIATIONS

BSA: Bovine Serum Albumine
ICG: indocyanine green
MSD: mean square displacement
NC: nanoclusters
NIR: near-infrared
PL: photoluminescence
QY: quantum yield
SPT: single particle tracking
SAXS: small angle x-ray scattering:
siHSF1: silencing heat shock transcription factor 1
TEM: transmission electron microscopy.

## Notes

### Competing Interest Statement

The authors have declared no competing interest.

## REFERENCES

(1) Chen, L., Black, A., Parak, W. J., Klinke, C., Chakraborty, I. Metal nanocluster-based devices: Challenges and opportunities. Aggregate 2022, 3 (4), e132. DOI: 10.1002/agt2.132 (acccessed 2025/01/20).

(2) Sahoo, S. R., Dinda, T. K., Saha, S., Mal, P., Goswami, N. Maneuvering the Electronic State and Active Site of Assembled-Gold Nanoclusters through Polyoxometalate Implantation for Heterogeneous Green-Light Photocatalysis. ACS Applied Materials & Interfaces 2025, 17 (13), 19669–19681. DOI: 10.1021/acsami.4c23033.

(3) Le Guével, X., Wegner, K. D., Würth, C., Baulin, V. A., Musnier, B., Josserand, V., Resch-Genger, U., Coll, J.-l. Tailoring the SWIR emission of gold nanoclusters by surface ligand rigidification and their application in 3D bioimaging. Chemical Communications 2022, 58 (18), 2967-2970, 10.1039/D1CC07055J. DOI: 10.1039/D1CC07055J.

(4) Casteleiro, B., Da Cruz-Boisson, F., Alcouffe, P., Pinto, S. N., Gaspar Martinho, J. M., Charreyre, M.-T., Farinha, J. P. S., Favier, A. Near-Infrared-Luminescent Gold Nanoclusters Stabilized by End-Grafted Biocompatible Polymer Chains for Bioimaging. ACS Applied Nano Materials 2023, 6 (13), 11689–11698. DOI: 10.1021/acsanm.3c01665.

(5) Santhoshkumar, S., Madhu, M., Tseng, W.-B., Tseng, W.-L. Gold nanocluster-based fluorescent sensors for in vitro and in vivo ratiometric imaging of biomolecules. Physical Chemistry Chemical Physics 2023, 25 (33), 21787–21801, 10.1039/D3CP02714G. DOI: 10.1039/D3CP02714G.

(6) van de Looij, S. M., Hebels, E. R., Viola, M., Hembury, M., Oliveira, S., Vermonden, T. Gold Nanoclusters: Imaging, Therapy, and Theranostic Roles in Biomedical Applications. Bioconjugate Chemistry 2022, 33 (1), 4–23. DOI: 10.1021/acs.bioconjchem.1c00475. Bonačić-Koutecký, V., Le Guével, X., Antoine, R. Engineering Liganded Gold Nanoclusters as Efficient Theranostic Agents for Cancer Applications. ChemBioChem 2023, 24 (4), e202200524. DOI: 10.1002/cbic.202200524 (acccessed 2024/10/03).

(7) Wu, Z., Du, Y., Liu, J., Yao, Q., Chen, T., Cao, Y., Zhang, H., Xie, J. Aurophilic Interactions in the Self-Assembly of Gold Nanoclusters into Nanoribbons with Enhanced Luminescence. Angewandte Chemie International Edition 2019, 58 (24), 8139–8144. DOI: 10.1002/anie.201903584 (acccessed 2025/03/26). Bonacchi, S., Antonello, S., Dainese, T., Maran, F. Atomically Precise Metal Nanoclusters: Novel Building Blocks for Hierarchical Structures. Chemistry – A European Journal 2021, 27 (1), 30–38. DOI: 10.1002/chem.202003155 (acccessed 2025/03/26).

(8) Compel, W. S., Wong, O. A., Chen, X., Yi, C., Geiss, R., Häkkinen, H., Knappenberger, K. L., Jr., Ackerson, C. J. Dynamic Diglyme-Mediated Self-Assembly of Gold Nanoclusters. ACS Nano 2015, 9 (12), 11690–11698. DOI: 10.1021/acsnano.5b02850.

(9) Yahia-Ammar, A., Sierra, D., Mérola, F., Hildebrandt, N., Le Guével, X. Self-Assembled Gold Nanoclusters for Bright Fluorescence Imaging and Enhanced Drug Delivery. ACS Nano 2016, 10 (2), 2591-2599, Article. DOI: 10.1021/acsnano.5b07596Scopus.

(10) Nie, Y., Tao, X., Zhang, H., Chai, Y.-q., Yuan, R. Self-Assembly of Gold Nanoclusters into a Metal–Organic Framework with Efficient Electrochemiluminescence and Their Application for Sensitive Detection of Rutin. Analytical Chemistry 2021, 93 (7), 3445–3451. DOI: 10.1021/acs.analchem.0c04682.

(11) Moro, S., Omrani, M., Erbek, S., Jourdan, M., Vandekerckhove, C. I., Nogier, C., Vanwonterghem, L., Molina, M.-C., Bernadó, P., Thureau, A., et al. Self-Assembled Peptide-Gold Nanoclusters with SiRNA Targeting Telomeric Response to Enhance Radiosensitivity in Lung Cancer Cells. Small Science 2025, 5 (2), 2400156. DOI: 10.1002/smsc.202400156 (acccessed 2025/01/28).

(12) Jiang, Y., Cao, H., Deng, H., Guan, L., Langthasa, J., Colburg, D. R. C., Melemenidis, S., Cotton, R. M., Aleman, J., Wang, X.-J., et al. Gold-siRNA supraclusters enhance the anti-tumor immune response of stereotactic ablative radiotherapy at primary and metastatic tumors. Nature Biotechnology 2024. DOI: 10.1038/s41587-024-02448-0.

(13) Lei, Y., Tang, L., Xie, Y., Xianyu, Y., Zhang, L., Wang, P., Hamada, Y., Jiang, K., Zheng, W., Jiang, X. Gold nanoclusters-assisted delivery of NGF siRNA for effective treatment of pancreatic cancer. Nature Communications 2017, 8 (1), 15130. DOI: 10.1038/ncomms15130.

(14) Haye, L., Diriwari, P. I., Alhalabi, A., Gallavardin, T., Combes, A., Klymchenko, A. S., Hildebrandt, N., Le Guével, X., Reisch, A. Enhancing Near Infrared II Emission of Gold Nanoclusters via Encapsulation in Small Polymer Nanoparticles. Advanced Optical Materials 2022, n/a (/a), 2201474, https://doi.org/10.1002/adom.202201474. DOI: 10.1002/adom.202201474 (acccessed 2022/10/14).

(15) Chang, H., Karan, N. S., Shin, K., Bootharaju, M. S., Nah, S., Chae, S. I., Baek, W., Lee, S., Kim, J., Son, Y. J., et al. Highly Fluorescent Gold Cluster Assembly. Journal of the American Chemical Society 2021, 143 (1), 326–334. DOI: 10.1021/jacs.0c10907.

(16) Olesiak-Banska, J., Waszkielewicz, M., Obstarczyk, P., Samoc, M. Two-photon absorption and photoluminescence of colloidal gold nanoparticles and nanoclusters. Chemical Society Reviews 2019, 48 (15), 4087-4117, 10.1039/C8CS00849C. DOI: 10.1039/C8CS00849C.

(17) Rival, J. V., Mymoona, P., Lakshmi, K. M., Nonappa; Pradeep, T., Shibu, E. S. Self-Assembly of Precision Noble Metal Nanoclusters: Hierarchical Structural Complexity, Colloidal Superstructures, and Applications. Small 2021, 17 (27), 2005718. DOI: 10.1002/smll.202005718 (acccessed 2025/01/20). Kolay, S., Bain, D., Maity, S., Devi, A., Patra, A., Antoine, R. Self-Assembled Metal Nanoclusters: Driving Forces and Structural Correlation with Optical Properties. In Nanomaterials, 2022; Vol. 12. Chen, H., Zou, L., Hossain, E., Li, Y., Liu, S., Pu, Y., Mao, X. Functional structures assembled based on Au clusters with practical applications. Biomaterials Science 2024, 12 (17), 4283–4300, 10.1039/D4BM00455H. DOI: 10.1039/D4BM00455H. Li, H., Kang, X., Zhu, M. Superlattice Assembly for Empowering Metal Nanoclusters. Accounts of Chemical Research 2024, 57 (21), 3194–3205. DOI: 10.1021/acs.accounts.4c00521.

(18) Simon, F., Weiss, L. E., van Teeffelen, S. A guide to single-particle tracking. Nature Reviews Methods Primers 2024, 4 (1), 66. DOI: 10.1038/s43586-024-00341-3. Wang, Z., Wang, X., Zhang, Y., Xu, W., Han, X. Principles and Applications of Single Particle Tracking in Cell Research. Small 2021, 17 (11), 2005133. DOI: 10.1002/smll.202005133 (acccessed 2025/05/08).

(19) Lee, A., Simon, A. A., Boyreau, A., Allain-Courtois, N., Lambert, B., Pradère, J.-P., Saltel, F., Cognet, L. Identification of Early Stage Liver Fibrosis by Modifications in the Interstitial Space Diffusive Microenvironment Using Fluorescent Single-Walled Carbon Nanotubes. Nano Letters 2024, 24 (18), 5603–5609. DOI: 10.1021/acs.nanolett.4c00955. Godin, A. G., Varela, J. A., Gao, Z., Danné, N., Dupuis, J. P., Lounis, B., Groc, L., Cognet, L. Single-nanotube tracking reveals the nanoscale organization of the extracellular space in the live brain. Nature Nanotechnology 2017, 12 (3), 238–243. DOI: 10.1038/nnano.2016.248. Somen Nandi; Quentin Gresil; Benjamin P. Lambert; Finn L. Sebastian; Simon Settele; Ivo Calaresu; Juan Estaun-Panzano; Anna Lovisotto; Claire Mazzocco; Benjamin S. Flavel; et al. Ultrashort Carbon Nanotubes with Luminescent Color Centers are Bright NIR-II Nano-Emitters. arXiv 2025, 2501.08254v2. DOI: 10.48550/arXiv.2501.08254.

(20) Simon, A. A., Haye, L., Alhalabi, A., Gresil, Q., Muñoz, B. M., Mornet, S., Reisch, A., Le Guével, X., Cognet, L. Expanding the Palette of SWIR Emitting Nanoparticles Based on Au Nanoclusters for Single-Particle Tracking Microscopy. Advanced Science 2024, n/a (/a), 2309267. DOI: 10.1002/advs.202309267 (acccessed 2024/05/20).

(21) Wang, F., Zhong, Y., Bruns, O., Liang, Y., Dai, H. In vivo NIR-II fluorescence imaging for biology and medicine. Nature Photonics 2024, 18 (6), 535–547. DOI: 10.1038/s41566-024-01391-5. Yang, Y., Xie, Y., Zhang, F. Second near-infrared window fluorescence nanoprobes for deep-tissue in vivo multiplexed bioimaging. Advanced Drug Delivery Reviews 2023, 193, 114697. DOI: 10.1016/j.addr.2023.114697. Sharma, N., Mohammad, W., Le Guével, X., Shanavas, A. Gold Nanoclusters as High Resolution NIR-II Theranostic Agents. Chemical & Biomedical Imaging 2024, 2 (7), 462–480. DOI: 10.1021/cbmi.4c00021.

(22) Yin, C., Xi, L., Wang, X., Eapen, M., Kukreja, R. C. Silencing heat shock factor 1 by small interfering RNA abrogates heat shock-induced cardioprotection against ischemia–reperfusion injury in mice. Journal of Molecular and Cellular Cardiology 2005, 39 (4), 681–689. DOI: 10.1016/j.yjmcc.2005.06.005 (acccessed 2025/05/16).

(23) Kang, X., Chong, H., Zhu, M. Au25(SR)18: the captain of the great nanocluster ship. Nanoscale 2018, 10 (23), 10758-10834, 10.1039/C8NR02973C. DOI: 10.1039/C8NR02973C.

(24) Yuan, X., Goswami, N., Chen, W., Yao, Q., Xie, J. Insights into the effect of surface ligands on the optical properties of thiolated Au25 nanoclusters. Chemical Communications 2016, 52 (30), 5234-5237, 10.1039/C6CC00857G. DOI: 10.1039/C6CC00857G. Musnier, B., Wegner, K. D., Comby-Zerbino, C., Trouillet, V., Jourdan, M., Hausler, I., Antoine, R., Coll, J. L., Resch-Genger, U., Le Guevel, X. High photoluminescence of shortwave infrared-emitting anisotropic surface charged gold nanoclusters. Nanoscale 2019, 11 (25), 12092–12096. DOI: 10.1039/c9nr04120f.

(25) Mohammad, W., Wegner, K. D., Comby-Zerbino, C., Trouillet, V., Ogayar, M. P., Coll, J.-l., Marin, R., Garcia, D. J., Resch-Genger, U., Antoine, R., et al. Enhanced brightness of ultra-small gold nanoparticles in the second biological window through thiol ligand shell control. Journal of Materials Chemistry C 2023, 11 (42), 14714-14724, 10.1039/D3TC03021K. DOI: 10.1039/D3TC03021K.

(26) Zhou, M., Jin, R., Sfeir, M. Y., Chen, Y., Song, Y., Jin, R. Electron localization in rod-shaped triicosahedral gold nanocluster. Proceedings of the National Academy of Sciences 2017, 114 (24), E4697–E4705. DOI: 10.1073/pnas.1704699114 (acccessed 2023/02/16).

(27) Xu, Q., Kumar, S., Jin, S., Qian, H., Zhu, M., Jin, R. Chiral 38-Gold-Atom Nanoclusters: Synthesis and Chiroptical Properties. Small 2014, 10 (5), 1008–1014. DOI: 10.1002/smll.201302279 (acccessed 2025/03/27).

(28) Liu, Z., Luo, L., Jin, R. Visible to NIR-II Photoluminescence of Atomically Precise Gold Nanoclusters. Advanced Materials 2024, 36 (8), 2309073. DOI: 10.1002/adma.202309073 (acccessed 2025/03/27).

(29) Liu, H., Hong, G., Luo, Z., Chen, J., Chang, J., Gong, M., He, H., Yang, J., Yuan, X., Li, L., et al. Atomic-Precision Gold Clusters for NIR-II Imaging. Advanced Materials 2019, 31 (46), 1901015, https://doi.org/10.1002/adma.201901015. DOI: 10.1002/adma.201901015 (acccessed 2023/02/16).

(30) Porret, E., Jourdan, M., Gennaro, B., Comby-Zerbino, C., Bertorelle, F., Trouillet, V., Qiu, X., Zoukimian, C., Boturyn, D., Hildebrandt, N., et al. Investigation of the Spatial Conformation of Charged Ligands on the Optical Properties of Gold Nanoclusters Journal of Physical Chemistry C 2019, 123 (43), 26705–26717.

(31) París Ogáyar, M., Ayed, Z., Josserand, V., Henry, M., Artiga, Á., Didonè, L., Granado, M., Serrano, A., Espinosa, A., Le Guével, X., et al. Luminescence Fingerprint of Intracellular NIR-II Gold Nanocluster Transformation: Implications for Sensing and Imaging. ACS Nano 2025, 19 (8), 7821–7834. DOI: 10.1021/acsnano.4c13955.

(32) Horita, J., Cole, D. R. Chapter 9 - Stable isotope partitioning in aqueous and hydrothermal systems to elevated temperatures. In Aqueous Systems at Elevated Temperatures and Pressures, Palmer, D. A., Fernández-Prini, R., Harvey, A. H. Eds., Academic Press, 2004; pp 277–319.

