## supporting information for "Self-Assembled siRNA-Gold Supraclusters Detected at the Single-Molecule Level in the NIR-II Window"

### I- EXPERIMENTAL SECTION

#### Materials

Tetrachloroauric(III) acid trihydrate($HAuCl_4 \cdot 3H_2O$), 4-mercaptobenzoic acid(*p*-MBA), Cysteamine, tributylamine, and ammonium acetate were obtained from Sigma-Aldrich; borane–trimethylamine complex was obtained from Alfa Aesar; and methanol and diethyl ether were obtained from VWR. Milli-Q water with a resistivity of 18.2 MΩ cm was used for all experiments.

All glasswares were cleaned with aqua-regia, rinsed with Milli-Q water, and dried before use.

#### Synthesis of AuCystA

The synthesis of AuCystA follows a similar approach to that used for $Au_{25}$(*p*-MBA)$_{18}$[1], In a round bottom flask Cysteamine (0.49 mmol, 38 mg) is first dissolved in 40 mL of methanol and 4 mL of water. Then, tetrachloroauric acid trihydrate ($HAuCl_4$,0.25 mmol, 100 mg) is added at an ambient condition and vigorous stirring, the solution gradually turns cloudy orange. After 20 minutes, 5 mL tributylamine (0.43 M) is added and left stirring for a further 10 minutes to form a gold thiolate/tributylamine complex.

Then, to induce a slow reduction of gold, 200 mg of trimethylamine borane is added under stirring for 2 h before adding another 200 mg. The solution is left under stirring overnight for 24 h to achieve the formation of the clusters.

**Purification**

Precipitation of the clusters is induced by adding 1 mL of 10% $NH_4OH$ solution and 30 mL of diethyl ether. After centrifugation, the supernatant which contains unwanted products is removed. Another cycle of dissolution/precipitation (1 mL $H_2O$; 5 mL of MeOH and 20 mL of $Et_2O$) is done.

Purification of the AuCystA cluster is done by selective precipitation. The precipitate is dissolved in 5 mL of water, and then 700 mg of ammonium acetate is added, followed by 20 mL of MeOH. Under these conditions, a precipitate appears which is separated by centrifugation. The supernatant contains a population of small clusters. Two cycles of dissolution/precipitation are done to improve purity of the cluster at the end, the last one using 1 mL $H_2O$; 5 mL of MeOH and 20 mL of $Et_2O$. At the end the precipitate is diluted in water and lyophilized to determine the concentration.

**Self-assembled AuCystA-siRNA**

Lyophilized AuCystA powder were dissolved in RNase free water to a concentration of 1mg/mL. AuCystA are then filtered at 0.2 microns. Then AuCystA were mixed with 0.25 µg of siRNA (corresponding to 17 pmol) solution at different AuCystA:siHSF1 weight ratios of 1:1, 2:1, 5:1, and 10:1. The solution was incubated for 30 min at room temperature to complete the binding of siRNA with AuCystA via electrostatic interactions.

We used siHSHF1 as siRNA chemically modified nucleotides synthesized by Eurogentec. Double-stranded HSF1 (heat shock factor 1) small interfering RNA (siRNA) sequences were designed as follows:

siHSF1: 5’-UAU GGA CUC CAC CUG GAU AA-TT-3’ (sense)

5’-UUA UCC AGG UGG AGU CCA UA-TT-3’ (antisense)

Double-stranded siRNA used contains two 2′-methoxy modifications at the positions 4 and 12 in both strands to improve siRNA stability, siRNA binding affinity with AuCystA, as well as to reduce the innate immune response. The two last phosphodiester linkage at 3′ ends were replaced by two phosphorothioates to protect siRNA from 3′-exonucleases in vitro and in vivo[2].

**Physico-chemical characterizations**

1. Mass spectrometry

The molecular weight of the AuCystA was determined by MALDI-tof in positive mode using an Autoflex Speed mass spectrometer (Bruker Daltonics) in linear positive mode. A saturated solution of α-cyano-4-hydroxycinnamic acid (HCCA) in TA30 (water/acetonitrile (70/30) with 0.1% trifluoroacetic acid) was used as the matrix. The sample was prepared by diluting the nanoclusters in water to a final concentration of 0.25–0.5 mg/mL, filtering with a ziptip (Merck Milipore), and adding the matrix at a ratio of 1:1 in water.

2. Electrophoretic Mobility shift assays

Electrophoretic mobility shift assay using 1% agarose gel containing TAE (Tris-acetate-EDTA) buffer at pH 7 was performed to evaluate the complexation between AuCystA and siRNA as well as the protection of siRNA against RNase degradation. Free siRNA (0.25µg) and AuCystA-siRNA complex at the weight ratio 5:1 were incubated or not at 37 °C with RNase A at different concentrations (1, 5, and 10 µg $mL^{-1}$) for 5 min. Samples were then incubated with RNase inhibitors (RNAsecure) at 60 °C for 15 min to inactivate RNase. After adding glycerol, each sample was loaded into the agarose gel for electrophoresis. Fluorescent AuCystA was detected in the SWIR window. In a second time, the resulting gel was stained with GelRed (Biotium, 41003) for RNA staining for 30 min at room temperature and siRNA was visualized on a Gel Doc imaging system (Bio-Rad).

3. Transmission electron microscopy (TEM)

TEM images of AuCystA and AuCystA-siRNA complex were determined by HRTEM JEOL2010 using a monochromated microscope working at 200 kV. Prior to imaging, the AuCystA and AuCystA-siRNA were dispersed on copper grids covered with a carbon film. Particle size in each condition was estimated on at least 100 particles using FIJI software. EDX measurements were performed to detect gold and phosphate elements within the self-assemblies.

4. DLS/ potential zeta

The hydrodynamic diameter and the zeta potential of the prepared nanomaterials dispersed in water and phosphate buffer (PBS 10 mM, pH4 – pH11) were measured in triplicate on a Zetasizer instrument from Malvern.

5. Small angle X-ray scattering (SAXS)

SAXS data were collected at the SWING beamline at SOLEIL synchrotron in France. A highly concentrated solution was prepared by slowly mixing 9 µL of siHSF1 (100 µM) with 18 µL of AuCystA (1 mg $mL^{-1}$) in RNase-free $H_2O$ in a final volume of 200 µL. Subsequently, 40 µL of the samples were directly injected into a 1.5 mm diameter and 10 µm thick-walled quartz capillary using a robotic arm equipped with electronic pipettes. The first sample is noted at $t_1$=10 min. Measurements from the same stock sample were made at 20 min, 30 min, 60 min, 120 min, 180 min, and 240 min. Buffer measurements were conducted before and after each sample injection. Samples were measured using the in-vacuum EigerX 4m detector with a distance of 6 m and a photon energy of 12 keV, enabling a $q$ range of $1.4 \times 10^{-3}$ to 0.185 $Å^{-1}$ (with $q = 4\pi\sin\theta/\lambda$).

Raw image processing, including masking and azimuthal averaging, was performed using the FOXTROT program. Buffer subtraction, curve averaging, and SAXS profile analyses were done with the ATSAS 3.2.1 software package, which includes AUTORG for calculating the radius of gyration ($R_g$) and GNOM for the calculation of pair-wise distance distribution, $P(r)$. Only the curve measured at 1/10 dilution, having enough signal-to-noise and not displaying interparticle interactions, is presented here.

6. Absorption and PL measurements

Absorbance measurements of AuCystA were done in 1 cm quartz cuvettes from Hellma GmbH at room temperature. The absorption spectra were recorded on an Evolution 201, Thermo Scientific UV-Vis spectrophotometer between 350 and 1000 nm. For other absorbance measurements 3 µL of each sample containing 30 ng and 150 ng of siHSF1 and AuCystA, respectively, were measured on a NanoDrop spectrophotometer.

Photoluminescence measurements were performed using a SWIR spectrometer (Wasatch). Samples were placed in a quartz cuvette and irradiated by a laser with excitation at $\lambda_{exc.}$ = 808 nm (120 mW/cm$^2$). The spectra were collected between the wavelengths of 880 nm and 1700 nm.

PLQY of the AuCystA was determined using fluorescence spectra of the reference ICG dye (Indocyanine green) and AuCystA in water, between 880 and 1500 nm, ($\lambda_{exc}$ = 808 nm). PLQY were calculated using the following formula.

$$QY = QY_{ref} \frac{A_{ref}}{A} \cdot \frac{\eta^2}{\eta_{ref}^2} \cdot \frac{I}{I_{ref}} \quad (1)$$

where, $QY_{ref}$ = quantum yield of the reference ICG dye (QY = 2.9%, $\lambda_{em}$ = 1000–1350 nm, in water)[16], $\eta$ = refractive index of $H_2O$ (1.333), $\eta_{ref}$ = refractive index of $H_2O$ (1.333), $I$ = integration of the emission spectra for the sample, $I_{ref}$ = integration of the emission spectra for the ICG dye, $A$ = optical density (0.06 OD at $\lambda_{exc}$ = 808 nm for the sample and 0.09 for the reference dye).

For photostability analysis, a 2mm cuvette was filled with 400 uL for each sample (AuCystA, AuCystA-siHSF1 and ICG) and excited using $\lambda_{exc}$ = 808 nm. The laser was left on for 20 minutes, during which pictures were taken every 2 minutes using a Nirvan 640ST camera (Princeton) in sub-SWIR windows using a long-pass (LP) filters of 1000 nm coupled to a 25 mm lens (Navitar).

The excitation source was an infrared laser ($\lambda_{exc}$ = 808 nm; 120 mW/cm$^2$). The images were then analyzed using ImageJ (Fiji), by measuring the intensity on a fixed area for each photo.

7. <u>Excitation- Photoluminescence maps:</u>

Excitation- Photoluminescence map measurements were performed in a 100 µL quartz cuvette from Hellma GmbH and recorded in NanoLog Spectrofluoremter (Horiba) at room temperature.

Excitation covers from 300 nm until 1000 nm with steps of 10 nm using a Xenon lamp, while the emission is recollected from 850 nm to 1600 nm with steps of 20 nm, with a integration time of 2 sec. When siRNA was introduced, measurements were performed 30 minutes after mixing, ensuring the well-formation of the assemblies. **(Figure 2e** and **Figure S15 b**)

8. NIR-II imaging (solution, and in capillaries)

NIR-II imaging was performed using a Princeton camera 640ST (900−1700 nm) coupled with a laser excitation source at λ = 808 nm (120 mW/cm2), using an architecture designed by Kaer Labs (electronics, acquisition and pre-processing software, and optical design for the excitation source). We used a short-pass excitation filter at 1000 nm (Thorlabs) and long-pass filters on the NIR-II camera at 1250 or 1400 nm (Thorlabs). A 25 mm or a 50 mm lens with an N.A. = 4 aperture (Navitar) was used to focus on the samples. Analyses were performed using FIJI software.

9. Epifluorescence widefield single molecule microscope:

A Nikon Eclipse Ti inverted microscope was equipped with a Plan APO IR 60×/1.27 water immersion objective (Nikon), along with a 488 nm laser (Obis Laser, Coherent) for sample illumination at 1.4 kW/cm². A 635 nm long-pass dichroic mirror (Di01-R635, Semrock) was placed in the cube to reflect and direct the laser to the sample, as well as a 650 nm long-pass emission filter (FELH 650, Thorlabs) to collect the fluorescence from the sample. As a detector, a low-noise SWIR InGaAs camera (C-RED 2, First Light Imaging) was employed.

**Sample preparation:** The sample preparation protocol is identical for all concentration conditions in the study of complexation. For AuCystA without siHSF1, AuCystA is diluted in RNase-free water (1:15 vol/vol), resulting in a final concentration of 30.5 µg/mL. For the conditions of different AuCystA: siRNA (w/w) ratios, the amount of siRNA is kept constant while progressively more concentrated AuCystA in RNase-free water solutions are added to achieve 3:1, 5:1, and 9:1 ratios. Notably, for the 3:1 ratio, the same AuCystA concentration as the no-siRNA sample was used to ensure accurate comparison. For single particle imaging, 1 μL of each solution is placed between two coverslips separated by an adhesive spacer having a well in its center. The coverslips were pre-treated with a plasma cleaner to prevent contamination and surface interactions with the sample. Unless otherwise specified, when siRNA was involved in the sample, measurements were conducted 30 minutes post-mixing to ensure the consistent formation of the assemblies.

10. Particle Size Determination via Diffusion Analysis:

Each sample was placed under the microscope and recorded with an exposure time of 33ms $(t_E)$. In **figure 3d**, for each time-point shown, two field of views (FOVs) were recorded for approximately 6000 frames (~3.5 minutes). **Figure 3c** presents data from 5 FOVs, each recorded for at least 7000 frames, while the supplementary figure includes one FOV recorded for more than 5000 frames.

The recorded movies were first analyzed using the software Fiji, where the super-localization of diffusing particles was performed using the ThunderSTORM plugin. Subsequently, the reconstruction of trajectories and their corresponding square displacement (MSD) was computed using a custom python script in which trackpy library is imported. Lastly, the diffusion coefficient (D) was determined by linear fitting of the ensemble MSD data (obtained from the number of detected trajectories indicated in each figure)[3] using the following equation :

$$MSD(t) = \varepsilon - \frac{4}{3} D t_E + 4Dt \qquad (2)$$

where the offset consists of two terms: the static localization uncertainty ($\sigma$) defined as $\varepsilon = 4\sigma^2$, and a negative term to take into account the effect of a finite camera exposure time $t_E$ when acquiring an image of a diffusive particle. Once the diffusion coefficient $D$ is calculated, we can determine the dynamic localization uncertainty by the offset of Eq (2) as well as the hydrodynamic particle radius ($r$) by the stokes-Einstein relation:

$$D = \frac{k_B T}{6\pi\eta r} \qquad (3)$$

where $k_B$ is the Boltzman coefficient, T the temperature and $\eta$ the dynamic viscosity.

To estimate the error of the calculated hydrodynamic radius, we took into consideration two factors: the slope precision of the linear fit (1 nm approx.) and the uncertainty related to the temperature which impacts the dynamic viscosity of the sample. Indeed, as sample temperature can be estimated between 19ºC and 25ºC, the dynamic viscosity ranges from 0.0010300 Pa·s to 0.00089274 Pa·s, both extreme cases were employed to calculated the error of the hydrodynamic particle radius. The major value retrieved for the radius is related to the maximum temperature considered (i.e 25ºC) together with the superior error of the slope fitting, whereas the minor value is related to inferior error of the slope when temperature is 19ºC. For the actual value of the hydrodynamic particle radius, the temperature considered was 22ºC, with a $\eta$ of 0.00095743 Pa·s.

### II- Supporting Figures

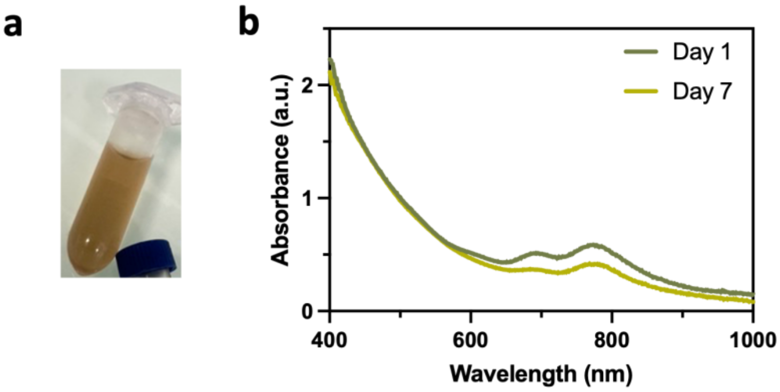


**Figure S1. (a)** Image of a Tube containing AuCystA in solution **(b)** Absorbance spectra showing stability of diluted AuCystA after 1 and 7 days from synthesis.

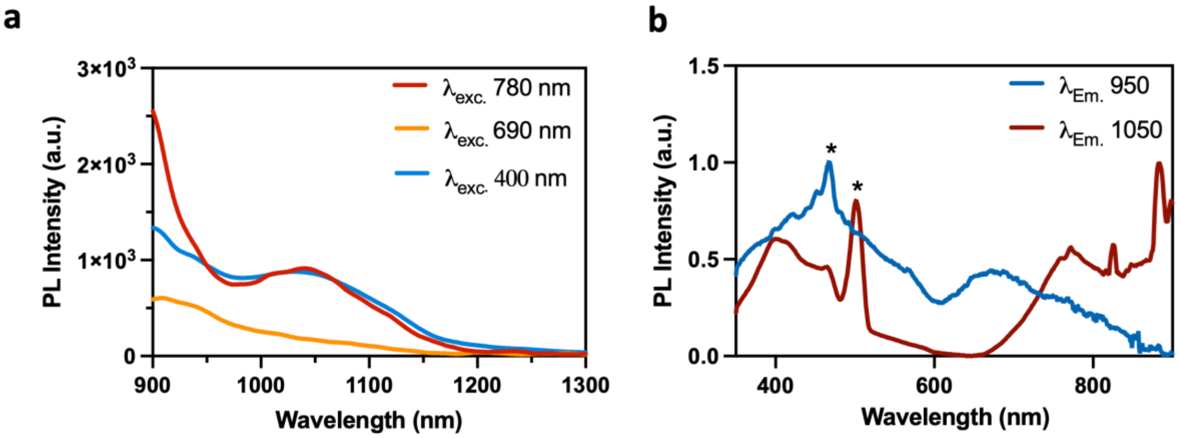


**Figure S2. (a)** PL Spectra of diluted AuCystA at different excitation wavelengths ($\lambda_{exc.}$ 400 nm, $\lambda_{exc.}$ 690 nm, and $\lambda_{exc.}$ 780 nm). **(b)** Excitation spectra of AuCystA with $\lambda_{em.}$ 950 nm and $\lambda_{em.}$ 1050 nm. *Artefact of the excitation laser.

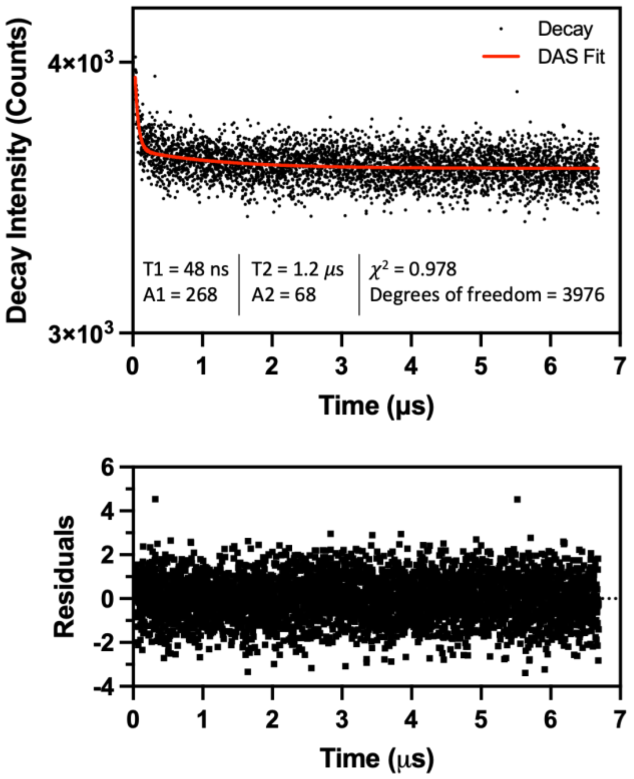


**Figure S3.** PL decay of AuCystA fit with biexponential decay function to obtain two PL lifetimes with their respective contributions with $\lambda_{exc.}$ 405 nm and $\lambda_{em.}$ 1050 +/- 20 nm.

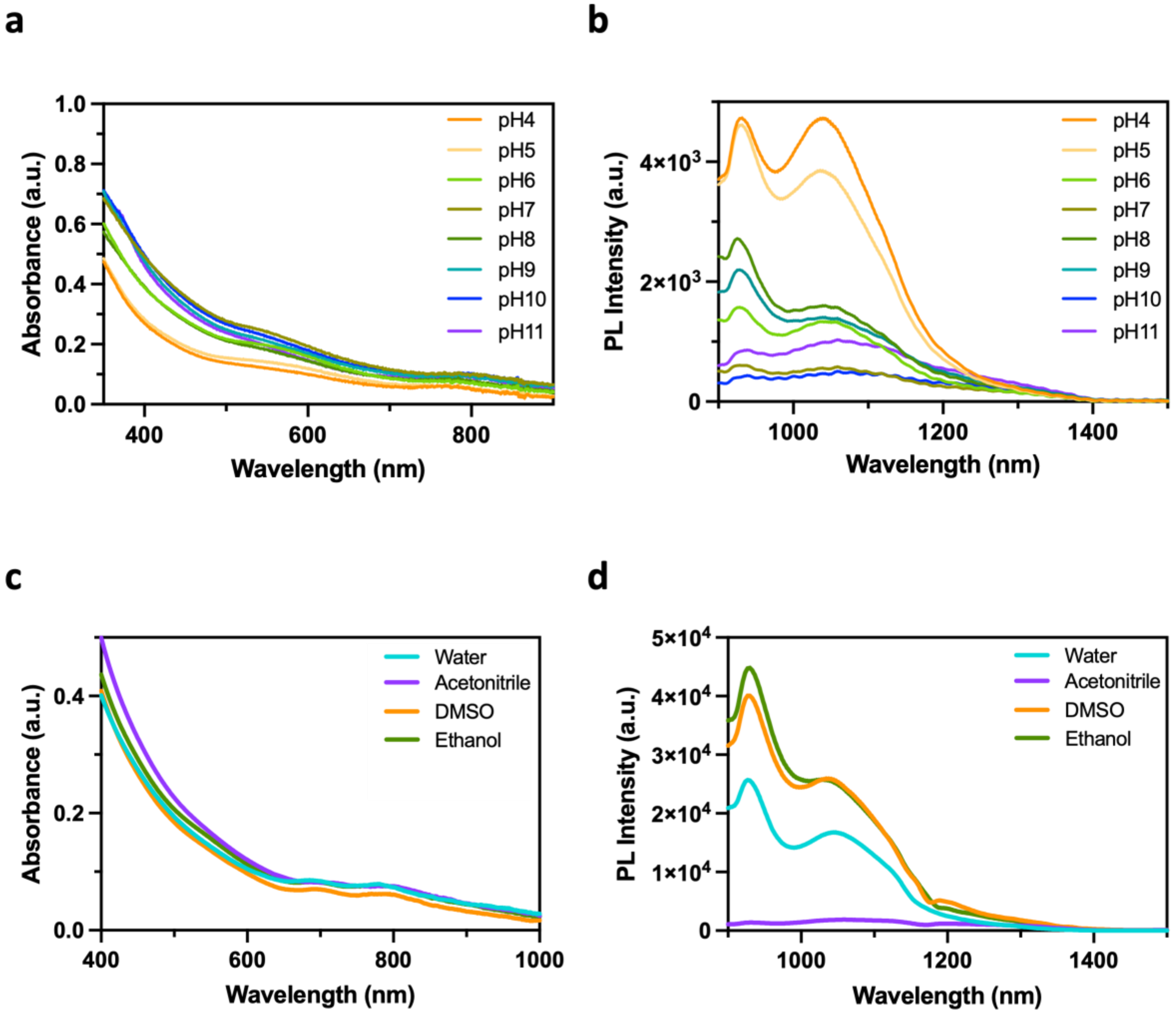


**Figure S4. pH and colloidal stability**

**(a)** Absorbance spectra of AuCystA diluted to the same concentration in PBS at different pH (Between pH4 to pH11). **(b)** PL spectra of AuCystA diluted to the same concentration in PBS at different pH (Between pH4 to pH11) $\lambda_{exc.}$ 808 nm. **(c)** Absorbance spectra of AuCystA diluted to the same concentration in different solvents (Water, Acetonitrile, Dimethyl sulfoxide (DMSO), and ethanol). **(d)** PL spectra of AuCystA diluted to the same concentration in different solvents (Water, Acetonitrile, DMSO, and ethanol) $\lambda_{exc.}$ 808 nm.

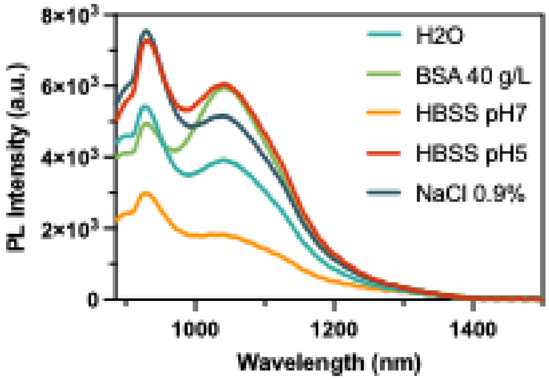


**Figure S5.** PL spectra of AuCystA in different biological buffer solutions. $\lambda_{exc.}$ 808 nm.

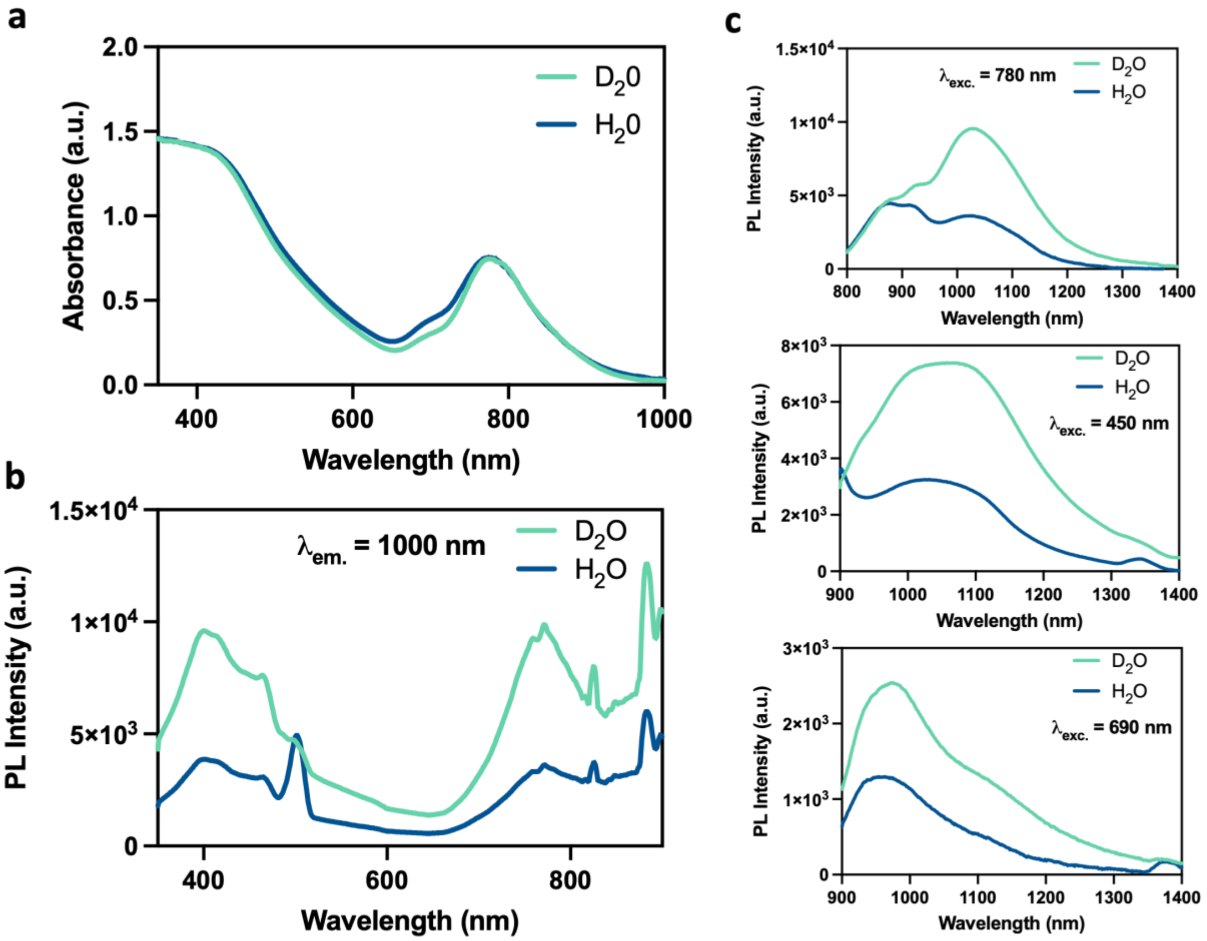


**Figure S6. (a)** Absorbance spectra of AuCystA (0.65 mg/mL) in $D_2O$ and $H_2O$. **(b)** Excitation spectra of AuCystA in $D_2O$ and $H_2O$ (0.65 mg/mL). $\lambda_{em.}$ 1000 nm. **(c)** PL Emission spectra of AuCystA (0.65 mg/mL) in $D_2O$ and $H_2O$ at $\lambda_{exc.}$ 780 nm, $\lambda_{exc.}$ 450 nm and $\lambda_{exc.}$ 690 nm.

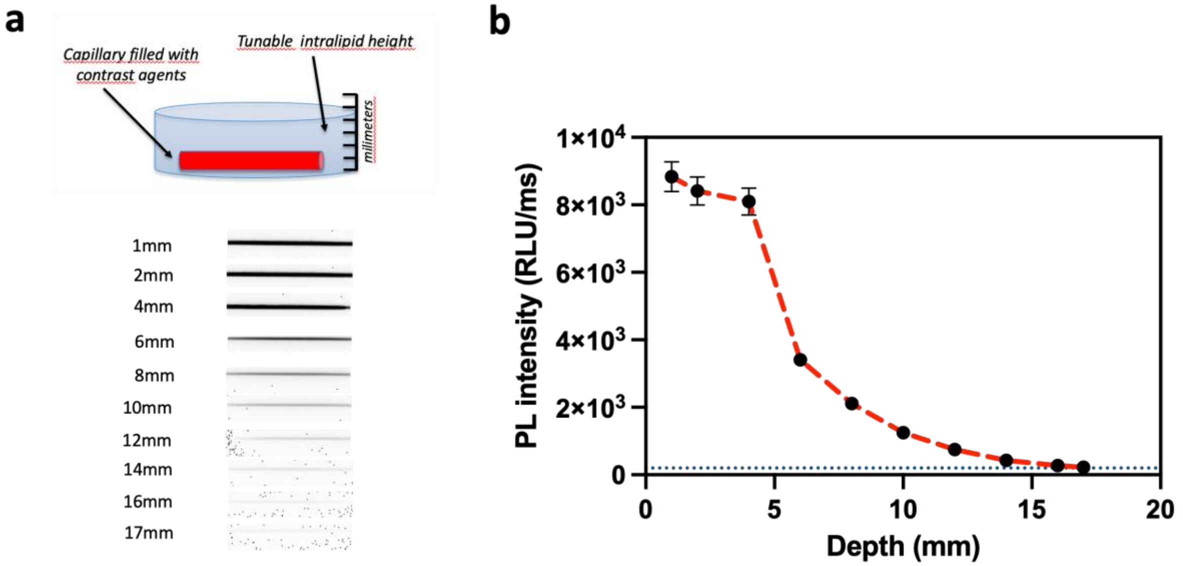


**Figure S7.** PL detection of AuCystA (1 mg/mL) in water in capillary soaked in intralipid solution (1%ww). $\lambda_{exc.}$ 808nm; $\lambda_{em.}$ 1250-1700nm. Exposure time: 100ms. Threshold (200 RLU/ms) of detection is shown by dotted line. **(a)** Schematic representation of the setup. **(b)** Graph extracted from the images showing the PL intensity in function of the depth.

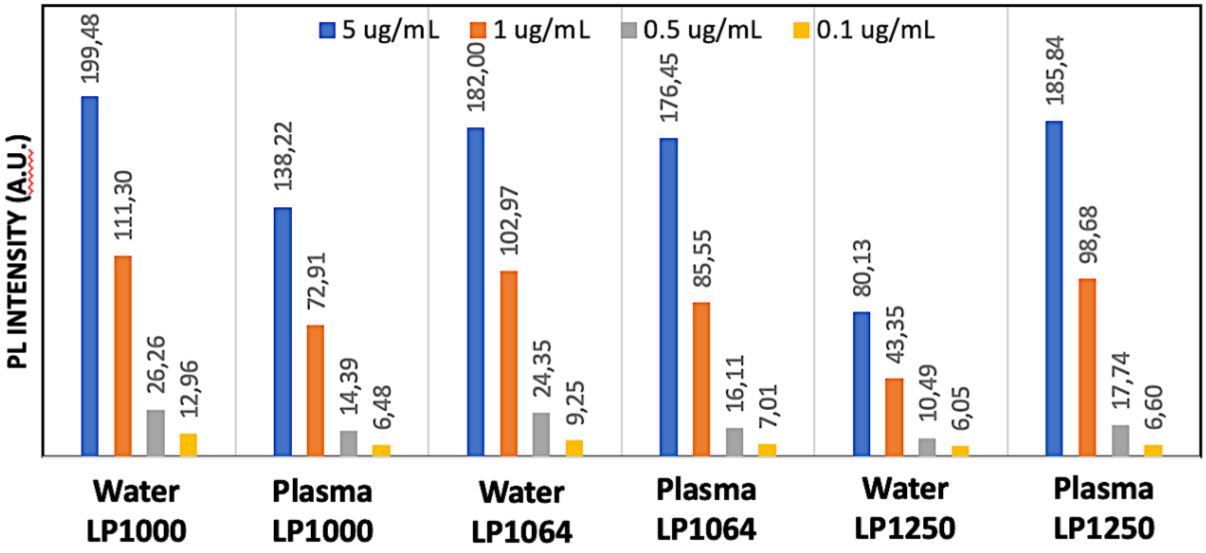


**Figure S8.** PL intensity of AuCystA at different concentration in water and in plasma in different sub-NIR-II windows. $\lambda_{exc.}$ 808 nm.

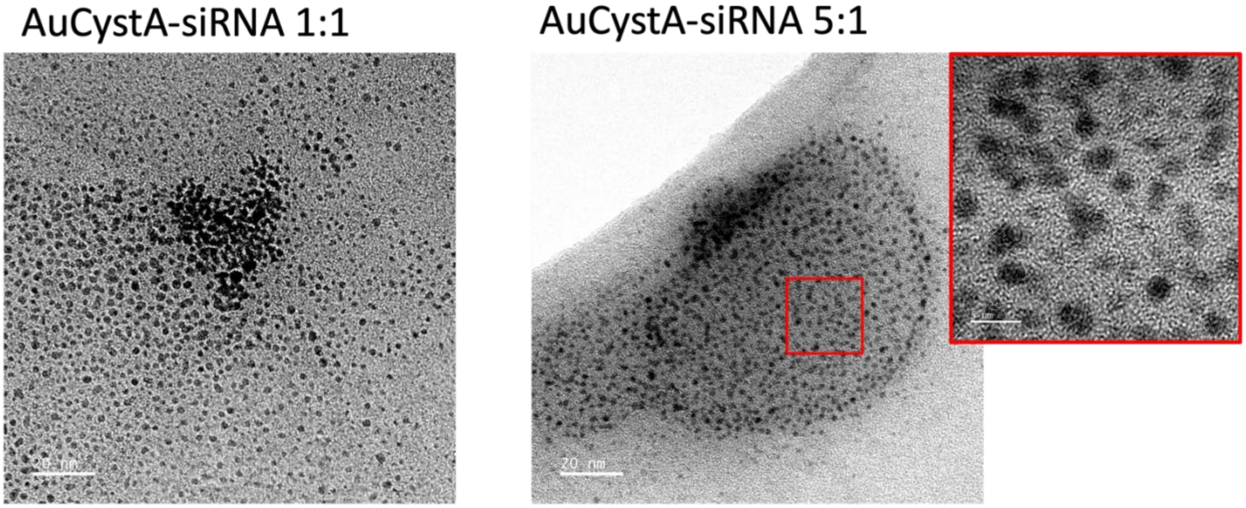


**Figure S9.** TEM images of the self-assembled AuCystA-siRNA at different weight ratios 1:1 and 5:1.

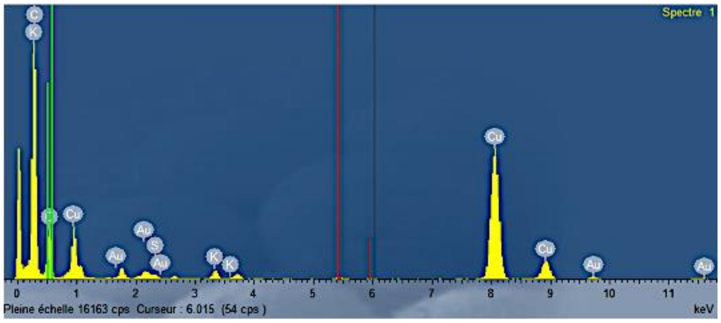


**Figure S10.** EDS Spectrum showing the different elements observed in the self-assembled AuCystA-siRNA (weight ratio 5:1).

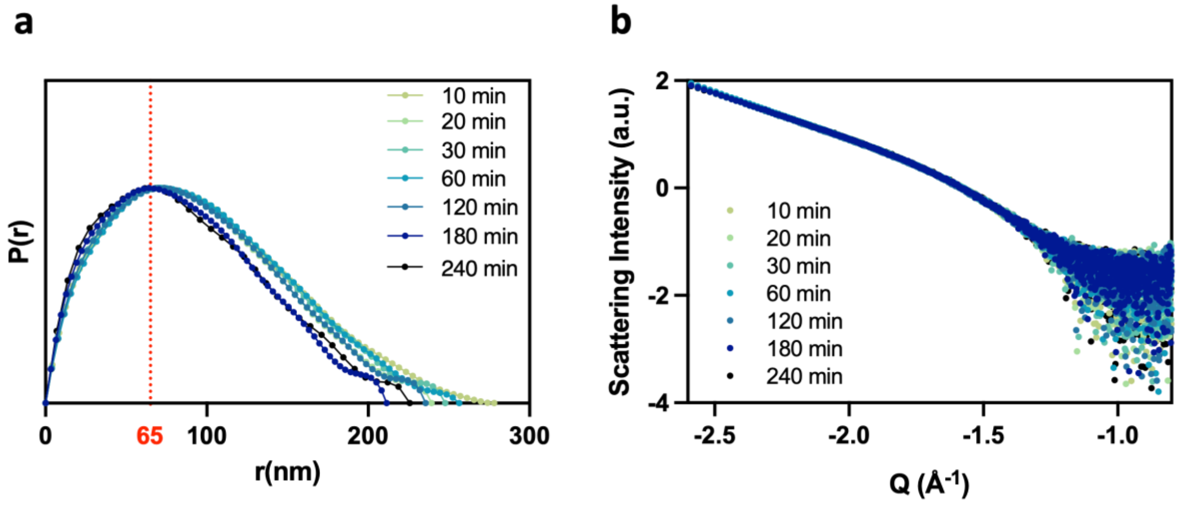


**Figure S11.** Pair-wise distance distribution P(r) (**a**), obtained from the Fourier transform of the SAXS curve (**b**) measured for a 1/10 dilution of the self-assembled AuCystA-siRNA prepared at the weight ratio 5:1.

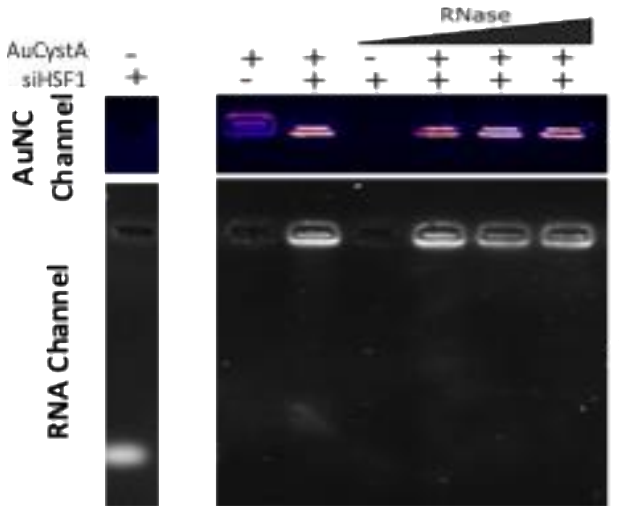


**Figure S12.** Migration of AuCystA and self-assembled AuCystA:siRNA (weigth ratio 5:1) in 1% agarose gel in the presence of increasing concentrations of RNase (1, 5, and 10 ng µL$^{-1}$).

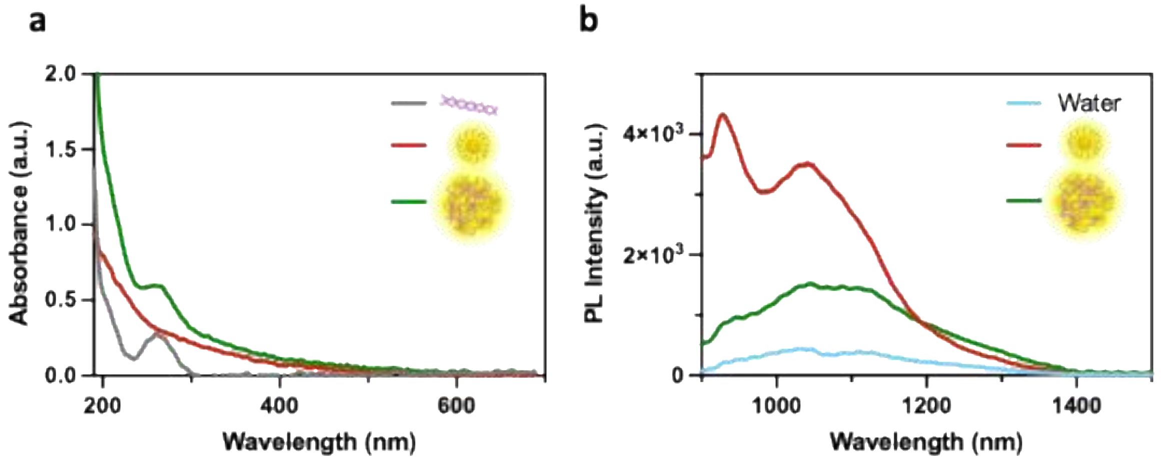


**Figure S13. (a)** Absorbance spectra of AuCystA at 50 µg/mL (red), siHSF1 at 10 µg/mL (grey) and self-assembled AuCystA:siRNA (weight ratio 5:1) at 50 µg/mL of AuCystA (green). **(b)** The respective PL spectra of AuCystA (red), Water (Light Blue) and self-assembled AuCystA-siRNA (green) with $\lambda_{exc.}$ 808 nm.

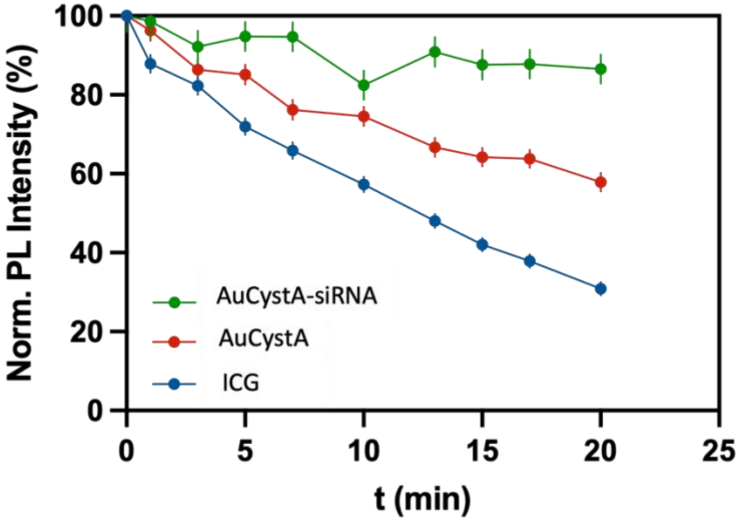


**Figure S14. Optical properties of AuCystA-siHSF1**

Photostability of AuCystA-siRNA (weigth ratio 5:1) (0.1 mg/mL ≈ 10 µM) and AuCystA (0.1 mg/mL ≈ 10 µM) compared to Indocyanine green (ICG) (10 µM). Measurements obtained from NIR-II imaging $\lambda_{exc.}$ 808 nm; $\lambda_{em.}$ 1050–1700 nm.

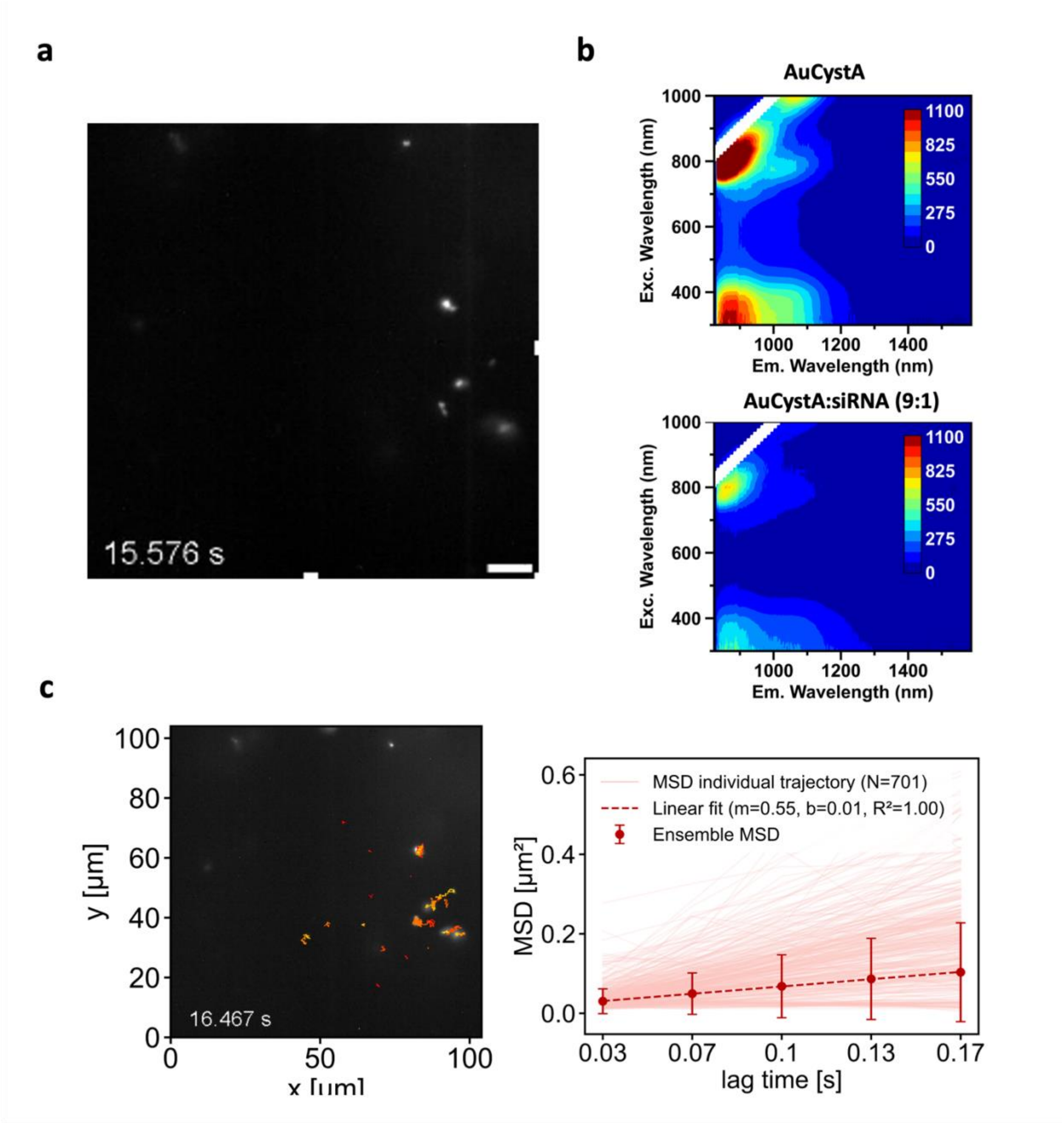


**Figure S15. NIR-II detection of AuCystA-siRNA at a weigth ratio 9:1.**

**(a)** Example image recorded after 16,467 s in the movie analysed in (c) and consisting of 502 frames (33 ms per frame). scale bar = 10 µm. **(b)** Excitation-photoluminescence maps. Left: AuCystA without siRNA; right: AuCystA–siRNA 9:1. A slight redshift is observed when siRNA is added, attributed to AuCystA aggregation. **(c)** MSD analysis of AuCystA–siRNA 9:1 (heterogenous structures). Individual MSDs (light red) from 701 trajectories in the same FOV; ensemble MSD calculated from individual MSD (dark red); linear fit of ensemble MSD (dark red dashed line) used to obtain the hydrodynamic radius of the tracked objects.

REFERENCES

(1) Bertorelle, F.; Russier-Antoine, I.; Comby-Zerbino, C.; Chirot, F.; Dugourd, P.; Brevet, P.-F.; Antoine, R. Isomeric Effect of Mercaptobenzoic Acids on the Synthesis, Stability, and Optical Properties of Au25(MBA)18 Nanoclusters. *ACS Omega* **2018**, *3* (11), 15635-15642. DOI: 10.1021/acsomega.8b02615.
(2) Chernikov, I. V.; Vlassov, V. V.; Chernolovskaya, E. L. Current Development of siRNA Bioconjugates: From Research to the Clinic. *Frontiers in Pharmacology* **2019**, *10*, Review. DOI: 10.3389/fphar.2019.00444.
(3) Michalet, X. Mean square displacement analysis of single-particle trajectories with localization error: Brownian motion in an isotropic medium. *Physical Review E* **2010**, *82* (4), 041914. DOI: 10.1103/PhysRevE.82.041914.
